# Auditory-Visual Interactions in the Blind with Artificial Vision: Are Multisensory Perceptions Restored After Decades of Blindness?

**DOI:** 10.1101/519850

**Authors:** Noelle R. B. Stiles, Vivek R. Patel, James D. Weiland

## Abstract

In the sighted, auditory and visual perception typically interact strongly and influence each other significantly. Blindness acquired in adulthood alters these multisensory pathways. During blindness, it has been shown that the senses functionally reorganize, enabling visual cortex to be recruited for auditory processing. It is yet unknown whether this reorganization is permanent, or whether auditory-visual interactions can be re-established in cases of partial visual recovery.

Retinal prostheses restore visual perception to the late blind and provide an opportunity to determine if these auditory-visual connections and interactions are still viable after years of plasticity and neglect. We tested Argus II retinal prosthesis patients (*N* = 7) for an auditory-visual illusion, the ventriloquist effect, in which the perceived location of an auditory stimulus is modified by the presence of a visual stimulus. Prosthetically-restored visual perception significantly modified patients’ auditory perceptions, comparable to results with sighted control participants (*N* = 10). Furthermore, the auditory-visual interaction strength in retinal prosthesis patients exhibited a significant partial anti-correlation with patient age, as well as a significant partial correlation with duration of prosthesis use.

These results indicate that auditory-visual interactions can be restored after decades of blindness, and that auditory-visual processing pathways and regions can be re-engaged. Furthermore, they indicate the resilience of multimodal interactions to plasticity during blindness, and that this plasticity can either be partially reversed or at least does not prevent auditory-visual interactions. Finally, this study provides hope for the restoration of sensory perception, complete with multisensory integration, even after years of visual deprivation.

**Significance:** Retinal prostheses restore visual perception to the blind by means of an implanted retinal stimulator wirelessly connected to a camera mounted on glasses. Individuals with prosthetic vision can locate and identify simple objects, and identify the direction of visual motion. A key question is whether this prosthetic vision will interact with the other senses, such as audition, in the same way that natural vision does. We found that artificial vision, like natural vision, can alter auditory localization. This suggests that the brain processes prosthetic vision similarly to natural vision despite altered visual processing in the retina. In addition, it implies that reorganization of the senses during blindness may be reversible, allowing for the rehabilitation of crossmodal interactions after visual restoration.

## Introduction

Over the past 20 years, sensory perception has been shown to consist not of isolated modalities but rather of richly interconnected senses that generate a holistic perception of the world (1, 2). In particular, studies on audition and vision have found that vision often dominates and modifies auditory perception, as illustrated by the McGurk and ventriloquist effects (3–5). Audition can also influence visual perception, especially when the stimuli occur over short time scales; illusions such as the double flash illusion and the flicker-flutter effect highlight this influence (6–10). These auditory-visual interactions allow the brain to improve decision making when sensory information is discordant or unreliable (11).

In parallel, a new consensus has developed that crossmodal plasticity occurs in the (late and early) blind, in which the viable senses (such as audition or somatosensation) re-purpose the visual cortex for non-visual sensory processing (12, 13). The visual cortex has been shown to be activated in the blind for auditory tasks such as sensory substitution perception, and for tactile tasks such as braille reading and roughness perception (14–18). This crossmodal plasticity likely plays a role in the heightened sensitivity in the blind for auditory localization and braille reading (19).

Nevertheless, it is as of yet unclear whether crossmodal plasticity during blindness will limit multimodal sensory integration in the visually rehabilitated. This is particularly relevant for the late blind, who have the highest potential for visual restoration as they had normal sensory perception and integration before losing visual perception. In addition, it is uncertain whether there is a re-learning period for the development of sensory integration in the visually restored, and whether the duration of blindness impacts the level of multimodality that can be restored. The development of retinal prostheses, which restore visual perception to the late blind, has provided a unique opportunity to investigate these questions.

The Argus II Retinal Prosthesis System restores visual function to those who became blind in adulthood with retinitis pigmentosa (RP) (20–24). The Argus II retinal prosthesis consists of a 60-element microelectrode array that electrically stimulates retinal cells based on visual information captured by a camera mounted on glasses (Fig. 1). Numerous studies have documented the visual experience of Argus II recipients, as well as other prostheses such as the Alpha-IMS (20–26). Visual perception with retinal prosthesis devices often begins with basic light perception, followed by the perception of visual flashes (phosphenes). With persistent rehabilitative training and device use, patients begin to see forms, and eventually can read letters and demonstrate improved navigation (25, 27). These patient outcomes have been reported to be relatively consistent between the two commercially available devices (the Argus II and Alpha-IMS) (25) and are characterized as ultra-low vision.

**Figure 1:**
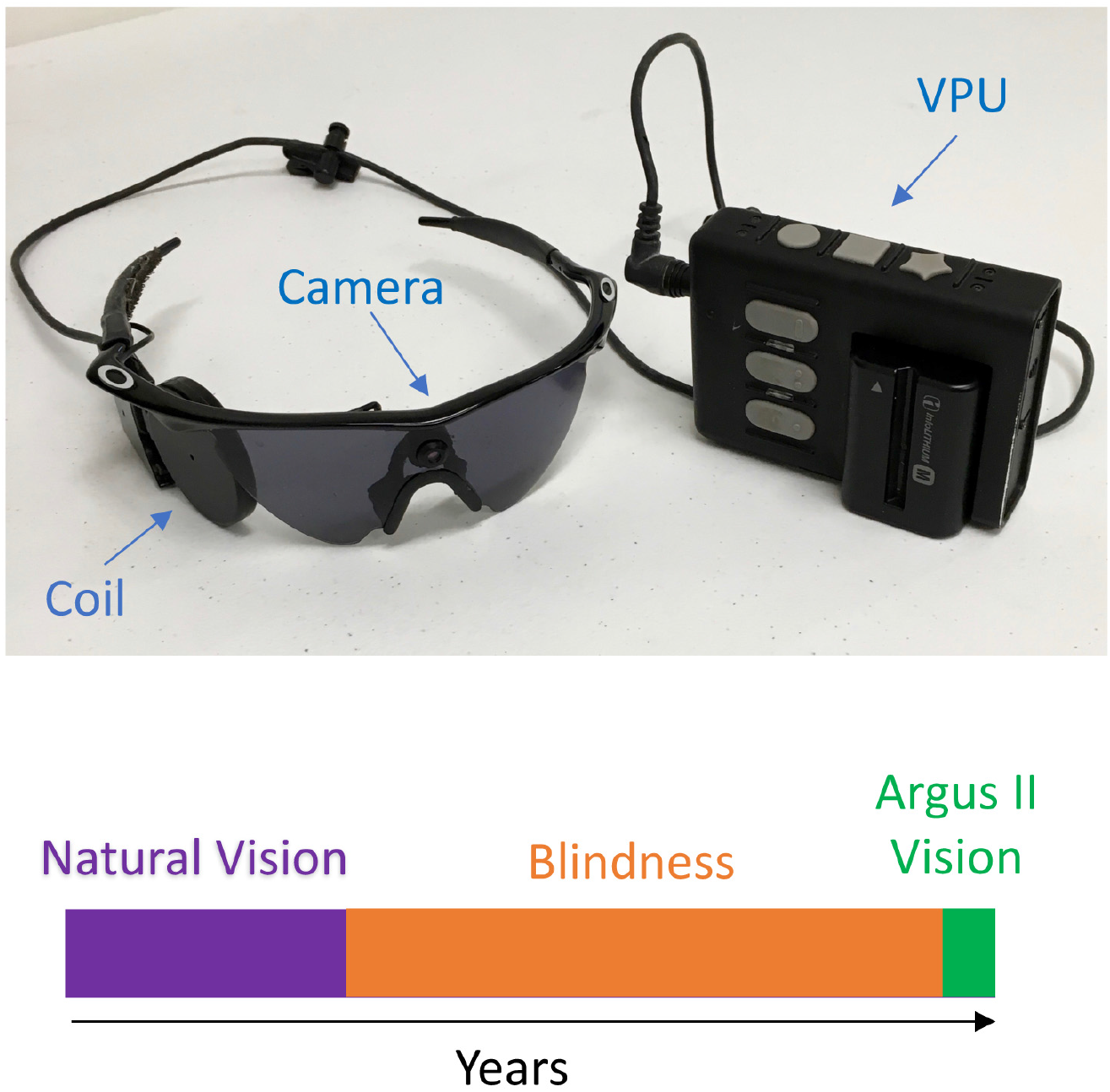
Argus II Retinal Prosthesis Device. An image of the Argus II external components is shown in the top of the figure, including the camera mounted on a pair of eyeglasses and the video processing unit (VPU). At the bottom of the figure is a diagram of the visual perception timeline of an Argus II patient: natural vision from birth, blindness in adulthood due to retinitis pigmentosa, and then restored perception with the Argus II device.

In this study, we show that this new artificial vision integrates with audition in a way similar to the multisensory integration of natural visual and auditory perception. In the ventriloquist effect, the perceived location of an auditory stimulus is modified by the presence of a visual stimulus (4–5). The Argus II patients (*N* = 7) demonstrated a significant auditory location shift when a paired auditory-visual stimulus was presented in comparison to the presentation of an auditory stimulus alone. Furthermore, the strength of this ventriloquist illusion in Argus II patients exhibited a significant partial anti-correlation with patient age, and a significant partial correlation with duration of prosthesis use.

## Results

### Ventriloquist Illusion Results in Argus II Patients (N = 7)

Seven late blind Argus II patients were tested for the ventriloquist effect 1.5 years or more after implantation with the retinal prosthesis device. Participants performed a localization task that was either unimodal (audition alone or vision alone), or bimodal (vision and audition together) (Fig. 2). For the visual task, participants used their Argus II prosthesis to localize a large white rectangular bar on a computer screen by moving their head-mounted camera to find it and then using a finger to point to it on the screen. The pointed position was then recorded by the experimenter by entering the closest reporting position (6.44 degrees between reporting positions, 9 positions on the screen). The auditory localization task required the subjects to listen to auditory beeps and then localize the sound by pointing to the screen surface where the beeps were perceived. Finally, the auditory-visual localization task was performed in two steps: (1) visual localization and pointing at a rectangular bar on the screen, then (2) auditory localization of beeps and pointing to their location (while still viewing the visual bar). During the second step (the auditory localization), the auditory stimulus and visual bar flashed and beeped in synchrony to promote binding (1).

**Figure 2:**
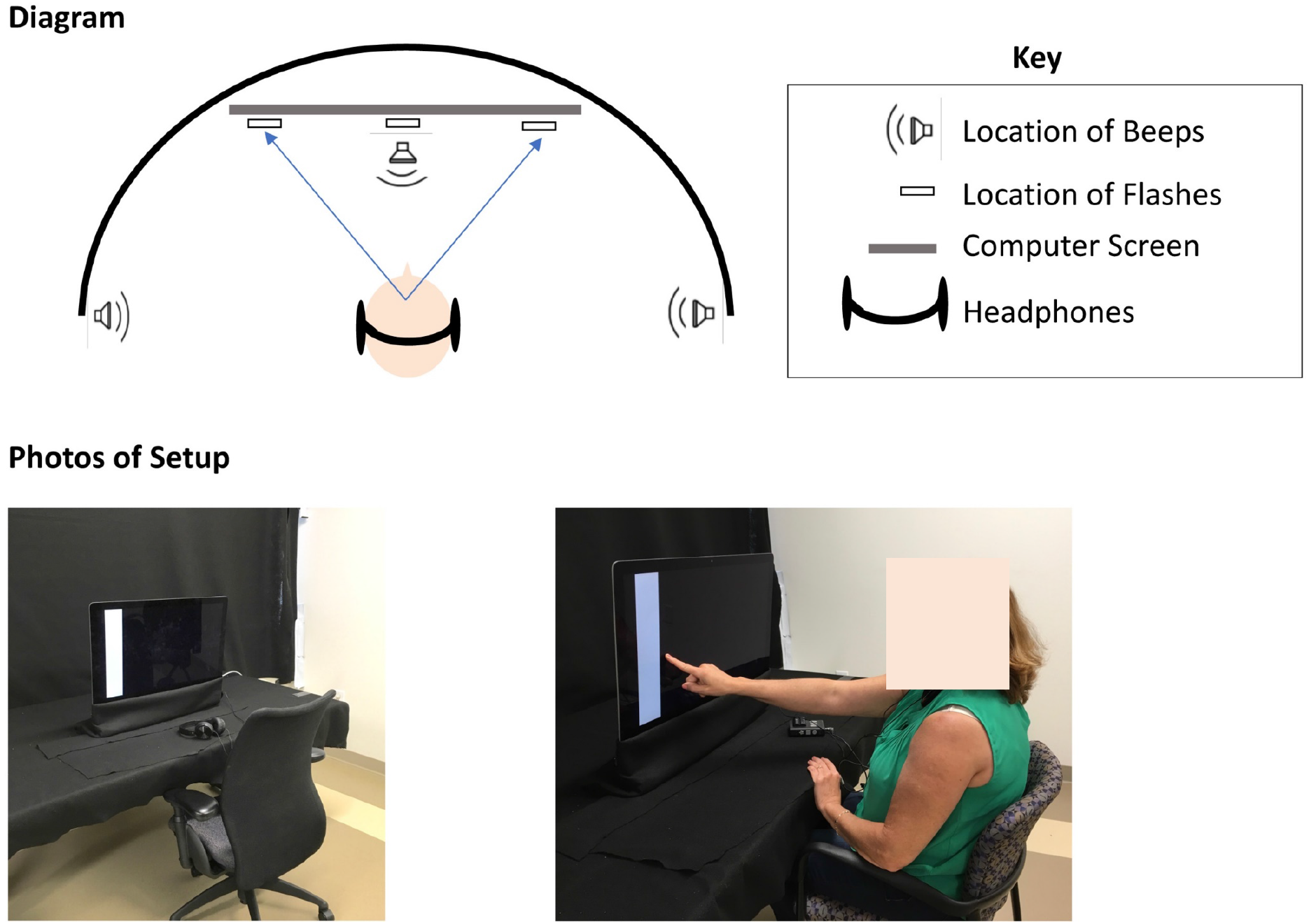
Experiment Setup. A diagram of the ventriloquist effect experimental setup is presented at the top of the figure. This diagram indicates the approximate locations of the auditory stimuli and the visual stimuli presented to the patients, and the extent of the screen where they reported auditory locations. At the bottom of the figure are two images of the experimental setup at the University of Southern California. The left image shows the experimental setup without a patient present, the right image shows the same experimental setup with a patient present. (Note: Video S1 demonstrates the performance of the ventriloquist task by an Argus II patient).

During the experiment, Argus II patients performed the unimodal tasks (vision alone and audition alone) to determine their baseline localization performance. The Argus II patients (*N* = 7) performed the spatial localization task with the visual bar alone or the auditory beep alone correctly (within one screen reporting position, 6.44 degrees between reporting positions, 9 positions on the screen) significantly above chance (Auditory (fraction correct): *M* = 0.74, *SD* = 0.13, *t*(6) = 12.84, *p* = 1.37 x 10^−5^; Visual (fraction correct): *M* = 0.71, *SD* = 0.08, *t*(6) = 19.11, *p* = 1.33 x 10^−6^) (Fig. S1). This indicates that the patients’ baseline visual perception and auditory perception were both sufficient for accurate localization.

To evaluate the magnitude of the ventriloquist effect, the reported auditory beep location (unimodal task, baseline) for each stimulus direction was subtracted from the corresponding reported auditory location during the auditory-visual trials (bimodal task). This shift of the auditory location (bimodal stimulus) relative to the auditory baseline (unimodal stimulus) was calculated for each participant in visual angle, such that a positive value indicates a shift toward the visual stimulus and a negative value indicates a shift away from the visual stimulus. This angular shift was significantly greater than zero (*M* = 5.80 degrees, *SD* = 3.34 degrees, *t*(6) = 4.59, *p* = 0.004) when averaged across all Argus II patients (*N* = 7) and all auditory locations (left, right, and center) (Fig. 3A).

**Figure 3:**
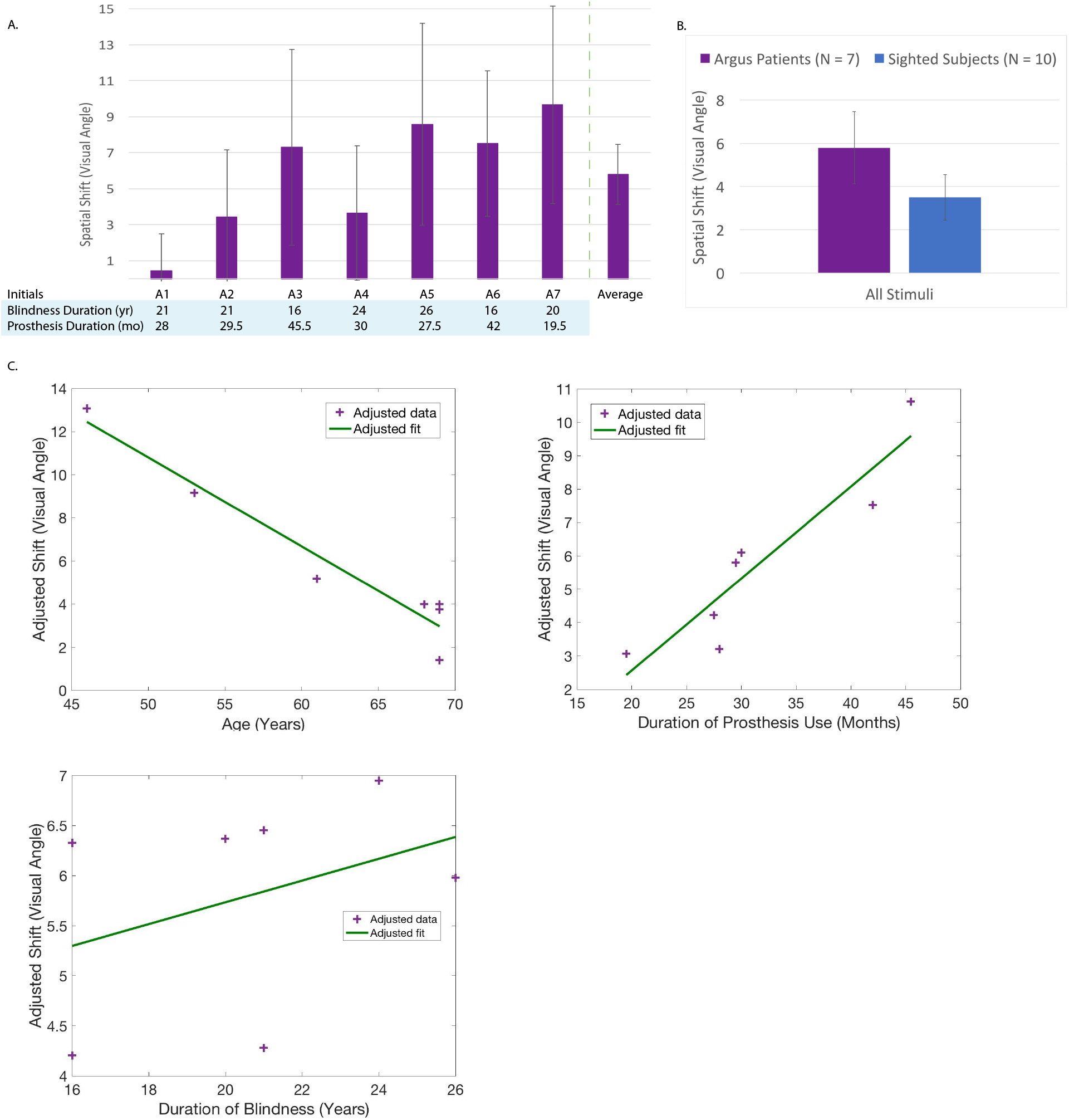
Ventriloquist Effect Results in Argus II Patients (*N* = 7) and Sighted Controls (*N* = 10) Panel A shows the size of the ventriloquist effect shift in the Argus II patients tested. The average shift of all of the patients is shown on the far right of the plot in Panel A with the bar titled “Average.” Panel B shows the comparison between the ventriloquist effect shifts of the Argus II patients (*N* = 7) and those of the sighted control participants (*N* = 10). Panel C of the figure shows the ventriloquist shift (in adjusted visual angle) as a function of age in years (left plot), the duration of prosthesis use in months (right plot), and the duration of blindness (bottom plot) (each data point represents one patient). The plots in Panel C are generated from a partial correlation, in order that the effect of each contributing factor of interest (age, duration of prosthesis use, and duration of blindness) on the ventriloquist shift can be isolated (see Methods for details). The *y*-values in the left plot of Panel C (*i.e*., the adjusted shift *vs*. age) are adjusted to take into account the duration of blindness and the duration of prosthesis use. The *y*-values in the right plot of Panel C (*i.e*., the adjusted shift *vs*. the duration of prosthesis use) are adjusted to take into account the duration of blindness and the age of the Argus II user. The *y*-values in the bottom plot of Panel C (*i.e*., adjusted shift *vs*. duration of blindness) are adjusted to take into account the duration of prosthesis use and the age of the Argus II user. Additional details on the methods of partial correlation and adjusted value plotting are under described in the Statistics subsection within the Methods section.

### Comparison Between Argus II Patients (N = 7) and Sighted Controls (N = 10)

Ten sighted control participants 50 years or older (*M* = 63.50 years, *SD* = 4.70 years) performed the same ventriloquist task completed by Argus II participants (*M* = 62.14 years, *SD* = 9.32 years). The sighted participants (*N* = 10) performed the spatial localization of the visual bar alone or auditory beep alone (unimodal tasks) correctly (within one screen reporting position, 6.44 degrees between reporting positions, 9 positions on the screen) significantly above chance (Auditory (fraction correct): *M* = 0.72, *SD* = 0.23, *t*(9) = 8.24, *p* = 1.75 x 10^−5^; Visual (fraction correct): *M* = 1.00, *SD* = 0, *p* = 0) (Fig. S1).

The sighted participants’ average ventriloquist shift (auditory localization with auditory-visual stimuli minus auditory localization with auditory alone stimuli) was significantly greater than zero (*M* = 3.64 degrees, *SD* = 2.15 degrees, *t*(9) = 5.36, *p* = 4.57 x 10^−4^). The sighted participants did not have a statistically significant difference in ventriloquist shift (*i.e*., influence of vision on auditory localization) relative to that of the Argus II participants (*t*(15) = 1.62, *p* = 0.13) (Fig. 3B and Fig. S2).

Nonetheless, it is perhaps counterintuitive that the Argus II participants appear to have a stronger (albeit not significantly different) ventriloquist shift than the sighted participants on average (5.80° ± 3.34° *vs*. 3.64° ± 2.15°). However, there are several reasons why this could be the case. First, the task performed was optimized for the ultra-low vision of the Argus II participants, and therefore the visual stimuli are repetitive and slow; as a consequence, many of the sighted participants commented that the task was monotonous. The ventriloquist effect has been found in other studies to be modulated by attention to the visual stimuli (28, 29), and therefore a lower degree of attention in the sighted participants may have weakened the effect. Second, the sighted participants experience other distracting and calibrating visual stimuli than just the visual stimuli on the screen. The sighted have a full view of the testing room with black felt, the experimenter’s face, walls, and the computer edges in their peripheral vision. This extra visual information is distracting from the stimuli, as well as adding additional visual fiducials that the auditory space is measured against. These additional distractors and fiducials could make the auditory localization less influenced by the visual stimulus on the screen. By way of contrast, the Argus II patients do not calibrate their auditory perception onto external visual space; they can only see the on-screen flash and none of the other 2D and 3D spatial markers. This lack of spatial calibration could make auditory localization more malleable in the Argus II participants. Finally, the Argus II patients likely had sensory plasticity during blindness, which altered the functional organization between auditory and visual cortical regions. This reorganization could have strengthened the connectivity between the sensory regions, thereby generating strong auditory-visual interactions.

### Correlation of Illusion Strength in Argus II Patients with Age and Duration of Prosthesis Use

We hypothesize that the age of the Argus II user, the duration of prosthesis use, and the duration of blindness could all potentially impact the degree of auditory-visual integration in Argus II users. Based on this hypothesis, a partial correlation analysis is the most appropriate statistical model to determine correlations between each of these factors and the ventriloquist shift. Using a partial correlation analysis, the age of the Argus II user (taking into account the duration of prosthesis use and the duration of blindness) was significantly anti-correlated with the auditory shift in Argus II patients (*N* = 7) (Fig. 3C) (*rho* = −0.95, *p* = 0.01). In addition, the duration of prosthesis use (taking into account the age of the Argus II user and the duration of blindness) was weakly correlated with the auditory spatial shift in Argus II patients (*N* = 7) (Fig. 3C) (*rho* = 0.83, *p* = 0.08). The duration of blindness before implantation of the retinal prosthesis was not significantly correlated with the auditory shift in Argus II patients (*N* = 7) (Fig. 3C) (*rho* = 0.27, *p* = 0.66).

When only the Argus II user age and duration of prosthesis use are used for a partial correlation, both factors correlate more strongly and significantly with the ventriloquist shift than when duration of blindness is also included in the linear model (Age of Argus II user: *rho* = −0.95, *p* = 0.004, Duration of Argus II use: *rho* = 0.86, *p* = 0.03).

These results indicate that the duration of blindness was not a critical factor in the ventriloquist effect modulation across these Argus II users. In particular, the patients in this study all have been blind for 15 years or more. Blindness for such a long time could mean that crossmodal reorganization due to vision loss will not be substantially altered with further years of blindness. Therefore, this result does not imply that the duration of blindness is not an important factor for those within a few years of the onset of blindness, but rather that it doesn’t play a critical role in differentiating patients with beyond a decade and half of blindness.

These partial correlation results also argue that the duration of prosthesis use and the age of the Argus II user are important indicators of the magnitude of auditory-visual interactions with Argus II ultra-low vision. This could mean that declining plasticity with age limits adaptation to prosthetic vision, and therefore hinders Argus II performance in the elderly. The positive correlation with duration of Argus II use could also indicate that a period of re-adaptation is required to re-learn visual processing and the relations between the senses. These possibilities will be explored further in the discussion section.

### The Role of Head Movement

The Argus II patients must scan the environment with their heads (as their visual input is provided by the Argus II head mounted camera) to locate and perceive visual stimuli. In our ventriloquist task, the participants moved their heads to locate the visual flash, and then kept their heads pointed toward the flash while they localized the auditory stimulus. Therefore, it is important to verify that head pointing toward the flash direction does not cause the shift in auditory localization, but rather that the influence of visual perception instigates this auditory shift. In particular, because the sounds are delivered via headphones, when the head is rotated it is possible that the perception of the sound is also shifted in the same direction. If this was the case, then a central flash would generate *less* of a shift (with peripheral sounds) than a flash on the left or right (with right or left peripheral sounds), as the head is not rotated when the flash is centrally located, while the head is rotated when the flash is on the left or right. However, the data indicate that a central flash (with peripheral sounds) has a larger shift than a flash on the left or right (with right or left peripheral sounds, respectively). Therefore, the shift toward the flash is not primarily due to head rotation.

It is also useful to note that Barneveld *et al*. studied the role of head and eye positions on the ventriloquist effect in sighted individuals and found that “the ventriloquist effect takes place in a common reference frame, rather than at a stage where audition and vision are still represented in their initial respective head-centered and eye-centered reference frames.” (30) This indicates that the crossmodal integration occurs *after* the transformations from the initial reference frames to a universal space, making head position not a determining factor on auditory positioning. While sighted participants have different head and eye integration than the Argus II patients, our results verify that head position was not a critical factor in generating the ventriloquist effect in this case.

### The Role of Cognitive Bias

Top down bias from prior knowledge, priming, other experimental cues, or confusion is always a concern with perceptual illusions. In particular, the ventriloquist effect could be influenced by cognitive biases, such as retinal prosthesis patients reporting the position of the visual flash when asked instead to locate the position of the auditory beep. If this were the case, then the measured auditory shifts toward the visual stimuli would tend to be increased. These types of reporting errors would be most likely to increase rather than decrease with increasing patient age. An argument against such cognitive bias with the ventriloquist illusion in this experiment is the observed partial anti-correlation between the illusion strength and Argus II patient age. It is further not likely that cognitive bias would increase with duration of Argus II use to generate the observed positive partial correlation between the illusion strength and Argus II duration of use.

To verify the existence of the ventriloquist effect in retinal prosthesis patients, the ventriloquist experiment was also performed using a double staircase method in two Argus II participants (A4 and A7) (based on the double staircase method of Berger and Ehrsson (31), and Bertelson and Aschersleben (32), as detailed below and in the supplemental information). The staircase ventriloquist task presents auditory stimuli on, for example, the far left, and asks the subject to identify the side from which the sound is perceived to originate (2AFC, left or right). If the subject identifies the side correctly, the sounds are then located closer to the center in the next trial, and if the side is incorrectly identified, the sounds are located more lateral (farther to the left or right) in the next trial. After a number of errors are made (between five and nine; see Supplemental Methods), the sound locations at which the errors were made are averaged to determine the threshold for discrimination of left-right sound location. In the double staircase version of the task, sounds are presented randomly on both the left and the right, and the left (negative) threshold is subtracted from the right (positive) threshold to derive a threshold difference (laterality index). This is equivalent to adding the magnitudes of the two thresholds.

Once the laterality index is obtained to identify the threshold for left-right disambiguation with the sound alone, the double staircase approach is used again with a central visual stimulus presented during the auditory task (visual distractor). If the ventriloquist effect is present, then the visual distractor should make the sound localization threshold more lateral (farther from the center) due to additional auditory localization error toward the center. For the two Argus II patients we tested, this method generated a significant difference in the audio alone threshold relative to the audio threshold with a visual distractor (Patient A4: *t*(8) = −4.43, *p* = 0.002; Patient A7: *t*(16) = −3.58, *p* = 0.003) (Fig. S5). This result represents a clear verification of the ventriloquist illusion. (Note: Additional experiments to verify that cognitive bias did not contribute to the ventriloquist illusion are described in the supplemental information.)

In conclusion, we verified that the ventriloquist illusion is not likely due to cognitive bias, based on the significant partial anti-correlation between the Argus II patients’ ventriloquist shift and their age, as well as with the double staircase ventriloquist experiment in two Argus II participants.

## Discussion

### Overview

This study investigated multimodal interactions in patients with partial vision restoration by means of retinal prostheses. The goal was to determine whether plasticity during blindness inhibits or limits crossmodal interactions when vision is restored in the late blind. We used the ventriloquist effect to test for integration between the artificial visual sense and the natural auditory sense in retinal prosthesis patients. Participants with implanted retinal prostheses exhibited a significant ventriloquist effect, in which prosthetic vision significantly modified auditory localization, that was not significantly different than that of sighted controls. In addition, it was found that the magnitude of the crossmodal influence partially anti-correlated with the age of the Argus II user, and partially correlated with the duration of prosthesis use. These results imply that crossmodal plasticity during blindness does not prohibit the restoration of multisensory interactions in the late blind. Furthermore, it indicates that a period of rehabilitation and learning is important for re-integration of the senses following prosthetic visual restoration.

### Implications for Auditory-Visual Interactions and Crossmodal Plasticity

Crossmodal plasticity has been shown to allow both auditory and tactile sensory processing to re-purpose the visual cortex in the blind (12, 13). Connections between sensory regions are reorganized and strengthened to allow for the capture of visual cortex for other processing priorities (12, 13). It has been unclear whether this extensive reorganization of cortical sensory regions fundamentally limits the potential for visual restoration, as well as the potential for crossmodal interactions following visual restoration. Very few individuals have recovered vision after blindness (early or late). As a consequence, whether crossmodal plasticity during late blindness limits either restored visual perception or crossmodal interactions following visual restoration has not been systematically tested; however, in the former case it has been hypothesized that it does (17).

Cochlear implants can restore audition to the deaf by means of electronic stimulation of the auditory nerve endings in the cochlea, and as such provide a useful comparison for the implications and progression of vision restoration by means of visual prostheses. It was found with the cochlear implant that the extent of cortical plasticity during deafness correlated with lower functionality following auditory restoration (33–35). These research findings implied that plasticity during sensory deprivation could reorganize auditory regions so extensively that they cannot be fully reconstituted, making the restoration of normal sensory processing limited.

Plasticity and reorganization during blindness could also limit the interactions between a restored sense and the other senses. Both feedback and feedforward networks for auditory-visual integration in the brain are known to be reorganized during blindness (12, 13), and therefore it is not clear if normal multisensory interactions can be restored. It was found in cochlear implant patients that they could integrate their restored audition with visual perception for lip reading (36–38). This result in cochlear implant patients provided a basis for us to hypothesize that this multimodal interaction could also be restored following blindness, where arguably the reorganization during blindness is potentially even more widespread than during deafness.

In the visual domain, the impact of brief visual deprivation on crossmodal integration during infancy has been previously investigated (39–43). The presence of dense cataracts for 5 to 24 months after birth was found to cause weaker (yet still present) crossmodal interactions years later when compared with normally sighted controls (40). In addition, brief cataracts at birth (up to 238 days) were found to enhance crossmodal activation (*i.e*., activation in visual cortex during an auditory task) later in life in comparison with that of normally sighted controls (39). While the visual deprivation period was brief for these patients, it occurred during the critical period, or the period from birth to adolescence when visual perception develops and is particularly sensitive to environmental changes (44). Therefore, this research indicates that early visual experiences are important to the development of crossmodal interactions.

The study described herein investigated the impact of prolonged visual deprivation in late blind patients (*i.e*., blindness onset in adulthood, after the critical period) on crossmodal interactions following partial (artificial) visual restoration. Our data support the claim that auditory-visual interactions can be restored in these patients, and that the interactions appear to strengthen with device use. These results bode well for the re-integration of the senses following blindness.

Although we found that auditory-visual interactions can be significantly restored in the late blind, the subjects in this study were all implanted following at least sixteen years of blindness. It is still unclear whether some of the inter-patient variability that we observe is attributable to differences in plasticity that were initiated following the onset of blindness. Additional experiments are required to determine if the extent of crossmodal plasticity during blindness (before prosthesis implantation) anti-correlates with the restoration of multimodal integration with a visual prosthesis. If this is true, then plasticity could limit but likely not eliminate the reintegration of the senses with visual restoration. This would provide critical insights into the impact of plasticity on the brain, and also indicate which patients have the greatest rehabilitative potential.

Finally, in patients with cochlear implants, plasticity following sensory restoration (*after* prosthesis implantation) has been harnessed to enhance sensory integration and perception, in some cases, even beyond that of individuals with natural hearing (37, 38). It will be interesting to determine if this type of enhanced multisensory integration can also occur with vision restoration. In particular, our comparison between sighted and Argus II patients indicates a potentially stronger ventriloquist effect in the Argus II patients. This difference could be due to alternative approaches to the task and different levels of attention, or could be due to the effects of crossmodal plasticity during blindness. Additional research comparing sighted and Argus II patients in crossmodal tasks is required to further clarify these potential differences.

### The Role of Age in Plasticity and Visual Rehabilitation

Our results indicate a partial anti-correlation between Argus II user age and auditory-visual interaction strength. This anti-correlation may be due to decreases in plasticity and adaptability in aging adults. Neuroimaging tools have been employed in gerontology research to study how and to what degree the brain can adapt and change in older adults. In particular, it has been shown directly with Transcranial Magnetic Stimulation (TMS) and Electroencephalography (EEG) that TMS-induced plasticity declines across the lifespan and may contribute to cognitive deficits in the elderly (45–47). In addition, magnetoencephalography (MEG) has been used to indicate that plasticity induced by a short training session also declines with age (48).

In line with these results, our experiments found that the older Argus II participants had less crossmodal interaction in the form of a weaker ventriloquist effect than younger participants. Crossmodal integration in Argus II patients depends on the plastic re-adaptation of visual cortex to interpret artificial visual input and to relate that input to existing auditory processing. We argue that the likely cause of the decline in crossmodal integration in Argus II users with age is based on the reduced capacity for plasticity in the elderly brain. However, additional experiments with more Argus II patients performing a wider variety tasks are needed to determine if age is a critical factor for predicting holistic rehabilitation outcomes with prosthetic vision.

### Implications for Argus Patient Rehabilitation

Auditory-visual interactions have been shown to be a critical component of sensory processing and behavior (1, 2). The key experimental result described herein that at least some multimodality can be restored following blindness is an important step toward the rehabilitation of the late blind with prostheses.

During blindness, patients have learned to rely on auditory and tactile perception to navigate their environment, to socialize with others, and to perceive their surroundings. When vision is restored, it is immeasurably important that the restored sense works synergistically with the existing senses. It is through these sensory interactions that the visually restored can potentially improve their daily lives while not relinquishing their other sensory skills.

In many ways, the sensory perception of the visually restored will not be the same as that of the naturally sighted. Reports of the visually restored have often conveyed more of a reliance on audition and tactile perception than the normally sighted (or reliance on vision in the case of cochlear implant patients) (38, 49). This may be due to crossmodal plasticity that was never reversed as well as personal routine and daily habits. Further experiments will be required to determine how best to integrate a restored sense with existing natural senses, and how to use crossmodal training paradigms to enhance the restored sense as well as performance in the tasks of daily living.

## Methods

### Participants

Seven participants implanted with the Argus II retinal prosthesis for 1.5 or more years took part in the experiment (3 Male, 4 Female). All of the participants had retinitis pigmentosa and at most light perception in one or both eyes. If they reported light perception in either eye, an opaque mask was used to occlude any natural visual perception within that eye (with the exception of one eye in one participant, detailed in Table S1). The duration of blindness (time following loss of vision up to the experiment) ranged from 16 years to 26 years, and participants’ ages at the time of the experiment ranged from 46 to 69 years of age (*M* = 62.14 years, *SD* = 9.32 years). One participant had an Argus I device implanted in one eye before receiving an Argus II device in the other eye. We tested only the Argus II device, and counted the years with the Argus I device as within the duration of blindness. Experiments with Argus II patients were performed at both the University of Michigan and the University of Southern California; in both locations the same researchers performed the experiment with the same equipment.

Ten sighted control participants performed the experiments at the University of Southern California (3 Male, 7 Female). Sighted participants were between the ages of 55 and 69 (*M* = 63.50 years, *SD* = 4.70 years), and a majority reported normal vision (with glasses or contacts) and no hearing loss (detailed in Table S2). Experiments with the sighted participants were performed using the same procedures as with the Argus II participants, including both oral instructions and the experimenter entering the participant responses in the computer.

All experiments were approved by either the University of Southern California or University of Michigan Institutional Review Boards, as appropriate, and all participants gave informed written consent.

### Experimental Tasks

The Three Sound Location Task methods and experimental design are detailed in the sections below. The other two tasks performed by a subset of participants (the Five Sound Location Task, and the Double Staircase Ventriloquist Task) are described in detail in the supplemental information.

### Stimuli (Three Sound Location Task)

Participants were seated such that the position of their eyes was about 20 inches away from an iMac 27-inch computer. The computer was placed on a black felt covered table, against a black felt covered wall, with the computer base and edge covered in black felt (Fig. 1). The room was dimly lit. Visual stimuli presented were vertical white rectangles (2.75 inches by 13 inches) on a black background presented for 0.5 seconds. The rectangles were presented on the far left, far right, or center of the computer screen. The auditory beeps presented were 0.07 seconds long (at 2,731 Hz) and delivered via headphones. The auditory location was indicated by the intensity difference between the left and right ear. The three locations were (1) far left (100% intensity left ear, 0% intensity right ear), (2) far right (0% intensity left ear, 100% intensity right ear), and (3) center (50% intensity left ear, 50% intensity right ear).

In the vision alone trials, the visual stimulus would flash on and off (0.5 seconds on, 0.5 seconds off) for 10.0 seconds (9.5 seconds from the beginning of the first flash to the end of the last of 10 flashes), and then return to the screen and remain stationary until the experimenter pressed the “1” key. After the “1” key was pressed, the visual stimulus would then flash 3 additional times (0.5 seconds on, 0.5 seconds off) (3.0 seconds total; 2.5 seconds from the beginning of the first flash to the end of the last of 3 flashes). In the auditory alone trials, the participants would hear a series of three auditory beeps (0.07 seconds on, 0.93 seconds off) for 2.07 seconds.

In the auditory-visual trials, the visual stimulus would flash on and off (0.5 seconds on, 0.5 seconds off) for 10.0 seconds (9.5 seconds from the beginning of the first flash to the end of the last of 10 flashes), and then remain stationary on the screen until the experimenter pressed the “1” key. Then the visual stimulus would flash on and off (0.5 seconds on, 0.5 seconds off) in synchrony with auditory beeps (each 0.07 seconds in duration, with the same onset as that of the flash) for 3.0 seconds (2.5 seconds from the beginning of the first flash to the end of the last of 3 flashes, in conjunction with 3 beeps).

### Task Description (Three Sound Location Task)

#### Overview

In the Three Sound Location Task, each stimulus type (auditory alone trials, visual alone trials, and auditory-visual trials) was grouped into a separate experiment sub-part. The experiment sub-parts were randomized in order and the trial order was also randomized within each sub-part. Each experiment sub-part had a different number of trials based on the number of stimulus locations (auditory alone trials, 3 auditory locations × 5 trials per stimulus location = 15 trials; visual alone trials, 3 visual locations × 5 trials per stimulus location = 15 trials; auditory-visual trials, 9 auditory-visual stimulus locations × 5 trials per stimulus location = 45 trials). Processed data presented in the Results section (the calculated auditory shift), and in Figs. 3A and 3B, only included trials with different auditory and visual locations (trials with the same location were excluded).

#### Auditory Alone Trials

Subjects were presented with auditory beeps on the left, right, or center for each trial. The stimulus order was randomized among the three locations. Subjects performed 5 trials for each location. At the beginning of each experiment, Argus II subjects were shown the edges of the screen with their hands. All participants were asked to point to the location on the screen of the auditory beep that they heard. The auditory beeps could be perceived within a range of 180 degrees, and the screen was about 60 degrees wide, and therefore the participants were asked to map the full auditory space onto the width of the screen. For example, a sound on the far left was mapped to the left edge of the screen, and so forth. The experimenter then entered the number closest to the location the participant pointed to within a range of 9 locations that were equally spaced on the computer screen.

#### Visual Alone Trials

Subjects were presented with visual flashes on the left, right, or center for each trial (the visual image remained on the screen until the flash was located and the experimenter pressed the “1” key). The stimulus order was randomized among the three locations. Subjects performed 5 trials for each location. At the beginning of each experiment, Argus II subjects were shown the edges of the screen with their hands. All participants were asked to point to the location of the visual flash that they perceived on the screen. The experimenter then entered the number closest to the location the participant pointed to within a range of 9 locations that were equally spaced on the computer screen.

#### Auditory-Visual Trials

Subjects were presented with visual flashes (on the left, right, or center) and asked to locate these flashes and point to them (the visual image remained on the screen until the flash was located and experimenter pressed the “1” key). Then the participant removed their pointed finger, but remained focused on the location of the visual flashes. Additional visual flashes with synchronous beeps were then presented, and the participant was asked to be aware of the flashes but to locate the beeps. The stimulus order was randomized among all visual and auditory stimulus combinations (9 combinations: 3 beep locations and 3 visual locations). Subjects performed 5 trials for each of the stimulus combinations.

At the beginning of each experiment, Argus II subjects were shown the edges of the screen with their hands. All participants were asked to point to the location on the screen of the auditory beep that they heard. The auditory beeps could be perceived within 180 degrees, and the screen was about 60 degrees wide, and therefore the participants were asked to map the full auditory space onto the width of the screen. The experimenter then entered the number closest to the location the participant pointed to within a range of 9 locations that were equally spaced on the computer screen.

### Statistics

We used two-tailed student t-tests for statistical significance calculations (MATLAB functions ttest and ttest2). Correlation analyses were performed with the partialcorri function in MATLAB. The plots shown in Fig. 3C were generated in MATLAB using the functions table, fitlm, and plotAdjustedResponse.

A partial correlation, based on a linear regression model, was used to evaluate the relationships between the ventriloquist shift and age, duration of prosthesis use, and duration of blindness. This particular model was used because we hypothesized that multiple factors (age, duration of prosthesis use, and duration of blindness) might contribute to the ventriloquist shift strength in each individual. In general, a partial correlation model is used to test for the relationship between two variables (dependent and independent variables), while controlling for the impact of additional independent variables on the dependent variable in question. The partial correlation values, rho and pvalue, were calculated with the partialcorri function in MATLAB, which uses a two-tailed Pearson correlation.

To generate the plots in Fig. 3C, we used the fitlm function to plot the adjusted responses (Fig. 3C) used in a partial correlation analysis. The linear model used by fitlm was *Y* = 1 + *X*1 + *X*2 + *X*3, in which *Y* = Ventriloquist shift, *X*1 = Duration of prosthesis use, *X*2 = Age of Argus II user, and *X*3 = Duration of blindness. The adjusted ventriloquist shifts plotted in Fig. 3C were generated using the plotAdjustedResponse function in MATLAB, which adds the residual to the adjusted fitted value for each data point.

### Duration of Prosthesis Use

The duration of prosthesis use was calculated to within a half-month using the following method. Patients were asked to report the date of Argus II implantation; this date was rounded to the beginning of the month, the end of the month, or mid-way through the month. If the day of implantation was not remembered exactly, the middle of the month was used. The number of months was then determined from this implantation date to the date of the experiment. The date of the experiment was also rounded to the beginning of the month, the end of the month, or mid-way through the month.

### Argus II Retinal Prosthesis System

The Argus II retinal prosthesis was developed by Second Sight Medical Products to restore limited resolution visual perception to advanced retinitis pigmentosa patients. The device consists of a camera mounted on the bridge of a pair of glasses, which communicates the video stream to a belt-worn Video Processing Unit (VPU) (Fig. 1, top). The VPU processes the raw video into stimulation parameters and transmits this information to an RF coil on the glasses (Fig. 1, top, coil). The visual information as well as power is sent from the external coil to the implant coil, and then relayed via cable to the microelectrode array tacked to the retinal surface. The microelectrode array has 6 by 10 microelectrodes for directly stimulating the inner surface of the retina (visual field of view of 11 by 19 degrees). Additional details about the prosthesis system are detailed by Zhou et al. (21).

Additional methods details are included in the supplemental information.

## Acknowledgements

We are grateful for support from the National Institutes of Health, the Philanthropic Educational Organization Scholar Award Program, and Arnold O. Beckman Postdoctoral Scholars Fellowship Program.

