## Supplemental Information for "Auditory-Visual Interactions in the Blind with Artificial Vision: Are Multisensory Perceptions Restored After Decades of Blindness?"

### Supplemental Results

#### Additional Analysis of the Three Sound Ventriloquist Task

The baseline performance of the participants in the Three Sound Ventriloquist Task, including both visual alone localization and auditory alone localization tasks, is summarized in Fig. S1. These localization tasks were used to determine individualized corrections for the auditory-visual ventriloquist task. Shown in the figure are the percentages of correct responses for both the Argus II retinal prosthesis patients and the sighted control subjects.

In Fig. S2, the auditory shift is plotted separately for two cases, the first in which the sounds were presented in either periphery (far left and far right) with the visual stimulus (flash) in the center, and the second in which the visual stimulus was presented in either periphery and the sounds were presented in the center. The auditory stimuli on the periphery extend beyond the visual screen width (Fig. 2, top), requiring the participant to remap 180 degrees of visual space into the width of the screen (60 degrees) for location reporting on the screen surface. This spatial remapping should generate smaller spatial shifts by a factor of three due to the compression of visual space (from 180 degrees to 60 degrees). In addition, while spatial remapping is easy for the sighted, it is a more challenging task for the blind, introducing additional error. For this reason, the auditory stimuli in the periphery were processed separately from the central auditory sounds (no spatial remapping) in Fig. S2. As expected, the central sounds displayed a significantly larger auditory spatial shift in comparison to the peripheral auditory sounds in Argus II patients (Center: $M = 14.08°$, $SD = 7.28°$, $t(6) = 5.11$, $p = 0.002$; Periphery: $M = 1.75°$, $SD = 1.85°$, $t(6) = 2.50$, $p = 0.05$; Significant Difference Center *vs.* Periphery: $t(12) = 4.34$, $p = 9.58 \times 10^{-4}$) (Fig. S2).

**Cognitive Bias: The Five Sound Ventriloquist Task**

To further verify that the ventriloquist effect was not due to cognitive bias in the sighted participants, a second experimental paradigm was tested in a subset of the participants (See Supplemental Methods section below). This experimental setup increased the auditory ambiguity by presenting five auditory locations (instead of three), and using the iMac internal speakers (instead of headphones) (Figs. S3 and S4). The auditory baseline percent correct substantially decreased for the five sound task relative to the three sound task in this subset of the sighted participants ($N = 3$) (three sound task (fraction correct): $M = 0.96$, $SD = 0.08$, Chance = 0.11; five sound task (fraction correct): $M = 0.36$, $SD = 6.80 \times 10^{-17}$, Chance = 0.18) (see Supplemental Methods for details). On the one hand, if the measured auditory shift is primarily due to crossmodal interactions, then this heightened auditory ambiguity should allow vision to more strongly influence auditory localization, resulting in an *increase* in the measured ventriloquist shift. If, on the other hand, the measured auditory shift is primarily due to cognitive bias, such as participants pointing to the visual stimulus instead of to the auditory stimulus (a cognitive error), then the increased number of auditory stimulus locations should result in a *decrease* in the average measured ventriloquist shift.

Sighted participants who performed both tasks did in fact demonstrate a significant increase in the ventriloquist shift for the five sound task relative to the three sound task (Fig. S3). Therefore, the five sound task verifies the presence of a strong ventriloquist illusion in the sighted participants. Furthermore, the five sound experiment also verifies that cognitive bias did not likely generate the ventriloquist illusion in this case. Rather, it indicates that the illusion is more likely due to crossmodal interactions, because as the auditory perception becomes more ambiguous (in the five sound task *vs.* the three sound task) vision influences auditory localization more strongly.

The five sound ventriloquist task was also tested in two Argus II participants (A4 and A7), to verify that cognitive bias did not generate their ventriloquist shift. For one participant the auditory localization baseline in the five sound task was less than chance, making the shift relative to that baseline invalid. The second participant exhibited a positive ventriloquist shift with the five sound task that was smaller than the three sound task ventriloquist shift. This represents a null result relative to our hypothesis of a larger shift with more auditory ambiguity, and could be due to a variety of reasons that include tiredness and reduced retinal excitability with continued prosthesis use (the three sound task was performed at the beginning of the experimental testing day, and the five sound task at the end of experimental testing day).

### Supplemental Methods

#### Methods for the Five Sound Task of the Ventriloquist Experiment

*Stimuli*

The auditory location was indicated by the intensity difference between the left and right speakers. The locations were: (1) far left (100% intensity left speaker, 0% intensity right speaker), (2) center-left (75% intensity left speaker, 25% intensity right speaker), (3) far right (0% intensity left speaker, 100% intensity right speaker), (4) center-right (25% intensity left speaker, 75% intensity right speaker), and (5) center (50% intensity left speaker, 50% intensity right speaker). The visual stimuli were the same as those used in the three sound task.

*Tasks*

Auditory Alone Trials

Subjects were presented with auditory beeps on the left, center-left, center, center-right, or right for each trial. The stimulus order was randomized among the five locations. Subjects performed 5 trials for each location. The auditory stimuli were presented using the built-in computer speakers (no headphones), so that the far left auditory stimulus location, for example, essentially coincided spatially with the left edge of the computer monitor. All participants were asked to point to the location on the computer screen surface from where they perceived the beep to have originated. The experimenter then entered the position closest to the location the participant pointed to (17 positions presented that were equally spaced on the computer screen, with 3.76 degrees between reporting positions).

Visual Alone Trials

Subjects were presented with visual flashes on the left, center, or right for each trial (the visual image remained on the screen until the flash was located and experimenter pressed the “1” key). The stimulus order was randomized among the three locations. Subjects performed 5 trials for each location. All participants were asked to point to the location on the computer screen surface where they perceived the visual flash. The experimenter then entered the position closest to the location the participant pointed to (17 positions presented that were equally spaced on the computer screen, with 3.76 degrees between reporting positions).

Auditory-Visual Trials

Subjects were presented with visual flashes (on the left, right, or center) and asked to locate these flashes and point to them (the visual image remained on the screen until the flash was located and experimenter pressed the “1” key). Then the participant withdrew their pointed finger, but remained focused on the screen. Additional visual flashes with synchronous beeps were then presented, and the participant was asked to be aware of the flashes but to locate the beeps. The stimulus order was randomized among all visual and auditory stimuli combinations (15 combinations: 5 beep locations, for 3 visual locations). Subjects performed 5 trials for each of the stimulus combinations. All participants were asked to point to the location on the computer screen surface from where they perceived the beep to have originated. The experimenter then entered the position closest to the location the participant pointed to (17 positions presented that were equally spaced on the computer screen, with 3.76 degrees between reporting positions).

*Percent Correct Calculations*

The percent correct for the five sound task was calculated by determining whether or not the participant pointed at one of the three reporting locations that were proximal to that of the sound stimulus position. For example, if the sound was located centrally, then the sound was positioned at the 9 position on the screen, and if the individual reported an 8, 9, or 10 location, their response was deemed to be correct. Whereas in the three sound task, a correct response required the participant to point within a 6.44° window around the actual sound location, in the five sound task a correct response required the participant to point within an 11.28° window around the actual sound location. The chance level for this reporting method was 3/17, as there were three potential correct answers out of 17 location options, as compared with a chance level of 1/9 in the case of the three sound task. The percent correct was calculated in this manner so that the chance level was higher for the five sound location task than the chance level for the three sound location task. The fact that the measured percent correct for the five sound task was observed to *decrease* relative to that of the three sound task provided a strong indication of the increased level of audio location ambiguity, as desired.

**Methods for the Double Staircase Ventriloquist Task**

*Stimuli*

Participants were seated about 20 inches away from an iMac 27-inch computer. The computer was placed on a black felt covered table, against a black felt covered wall, with the computer base and edge covered in black felt (Fig. 1). The room was dimly lit.

The same auditory stimulus was used in both the auditory alone double staircase and the visual distractor double staircase experiments. In both the auditory alone and visual distractor double staircase tasks, the auditory location was indicated by the intensity difference between the left and right ears. The locations ranged from far left (100% intensity left speaker, 0% intensity right speaker), to far right (0% intensity left speaker, 100% intensity right speaker) in a number of steps ($N_S$ = 100), with increments of 2% intensity between each step. The auditory stimuli were presented in headphones for both the auditory alone and the visual distractor double staircase tasks.

In the auditory only double staircase task, the auditory stimulus beeped 4 times for each trial, and the pause between beeps was 0.75 seconds long. In addition, the auditory beeps presented were 0.07 seconds long (at 2,731 Hz).

In the visual distractor double staircase task, both a visual stimulus and an auditory stimulus were presented together. The visual distractor stimulus was a vertical white rectangle (2.75 inches by 13 inches) in the center of a black background on the iMac computer screen. The visual distractor stimulus flashed 4 times in synchrony with the 4 auditory beeps (at 2,731 Hz). The visual distractor stimulus and auditory beeps were presented with the following timing: [beep (0.07 sec), then flash (0.25 seconds), then pause (0.5 seconds)]. The [timing sequence] was repeated three times (four total pairs of beeps and flashes). During the visual distractor task, the Argus II participants wore an eye mask to prevent perception with any remaining natural vision.

*Task*

The participant was told to identify whether the sounds were presented on the left or the right side (2AFC). In addition, during the visual distractor double staircase task, the participant was also told to pay attention to a visual stimulus on the screen in front of them while listening to the sound and responding as to its location (2AFC, left or right). The experimenter was seated next to the participant, and entered the participant's responses into a computer using a keyboard.

*Experimental Design*

The staircase method was used to determine the auditory location threshold for correct left-right discrimination. In this method, a staircase is defined as a set of stimuli starting from one extreme in which it is easy to perform the task, and then increasing in difficulty as the participant correctly responds to inquiries. In this case, the initial auditory stimuli were presented on the far left for the left staircase (100% amplitude in the left ear, and 0% amplitude in the right ear), and on the far right for the right staircase (0% amplitude in the left ear, and 100% amplitude in the right ear). In a given trial, if the sound location was correctly identified, the task difficulty was increased by moving the sound location towards the midline by one step. We defined one step as a 2% amplitude change in the sounds presented to each ear. Alternatively, if the sound was incorrectly localized, we decreased the task difficulty by moving the sound location towards the initial location (*i.e.*, further left for the left staircase and further right for the right staircase) by one step. If the sound presented to the participant is in the initial location (such as far left for the left staircase) and answers the question incorrectly, the same initial location is repeated in the next trial. The progression of trials in each staircase continues until the participant answers 9 trials incorrectly, or the participant performs 100 trials, whichever comes first.

A double staircase design performs the left and right staircases in the same block of trials by interleaving the trials for each staircase. In our experimental design, we randomly selected the staircase side to be tested in each trial, and went back and forth between the two staircase sides until either both staircases had 9 reversals each or the participant performed 100 trials total. One of the Argus II retinal prosthesis participants (A7) reached the 9 incorrect trial limit. The second participant (A4) reached 100 trials before reaching the 9 incorrect trial threshold, in particular reaching the 9 incorrect threshold on one side, but only 5 incorrect trials on the other. These incorrect answers are called reversals, and their location (fractional amplitude for the primary ear) is averaged to generate a threshold for that staircase, positive for the right staircase and negative for the left staircase. Therefore, the left and right staircase thresholds for A7 were calculated by averaging the locations of the first 9 reversals on each side, and the thresholds for A4 were calculated by averaging the locations of the first 5 reversals on each side. We then calculated the threshold difference (laterality index) in each case by subtracting the (negative) left staircase threshold from the (positive) right staircase threshold.

The double staircase task (left and right staircases) was performed twice by the participant, once with only auditory stimuli present, and a second time with a visual distractor presented on the screen accompanied by the auditory stimuli. The double staircase task was identical in both cases except for the presence of the visual distractor in the auditory-visual double staircase task. One of the participants (A4) was asked to verify that the visual stimulus was seen in each trial preceding the participants' auditory location response.

*Threshold Difference (Laterality Index) Calculation in Figure S5*

The threshold metric plotted in Fig. S5 was calculated by subtracting the thresholds (fractional amplitude in the primary ear) for the right staircase (positive value) and left staircase (negative value) using the same method as Berger and Ehrsson, Figs. 2C and 2D. The threshold difference with auditory stimuli alone was then compared to the threshold difference for the visual distractor stimulus double staircase (Fig. S5). The threshold differences were observed to be lower (threshold locations closer to the midline) when only auditory stimuli are present, and higher (threshold locations closer to the left and right edges) when the visual distractor is added to the auditory stimuli, as expected.

**Supplemental Videos**

**Video S1: The Auditory-Visual Ventriloquist Task**

This video shows a participant performing the ventriloquist task (auditory-visual trials only) that is detailed in this paper. The video has two parts; the first part has a demonstration of the ventriloquist task steps performed by a lab member, and the second part shows the task performed by an Argus II user. In Part 2, a subset of the auditory-visual trials that are typical of the average performance of this participant are shown.

**Supplemental Figures**

**Figure S1: Auditory and Visual Localization Percent Correct**

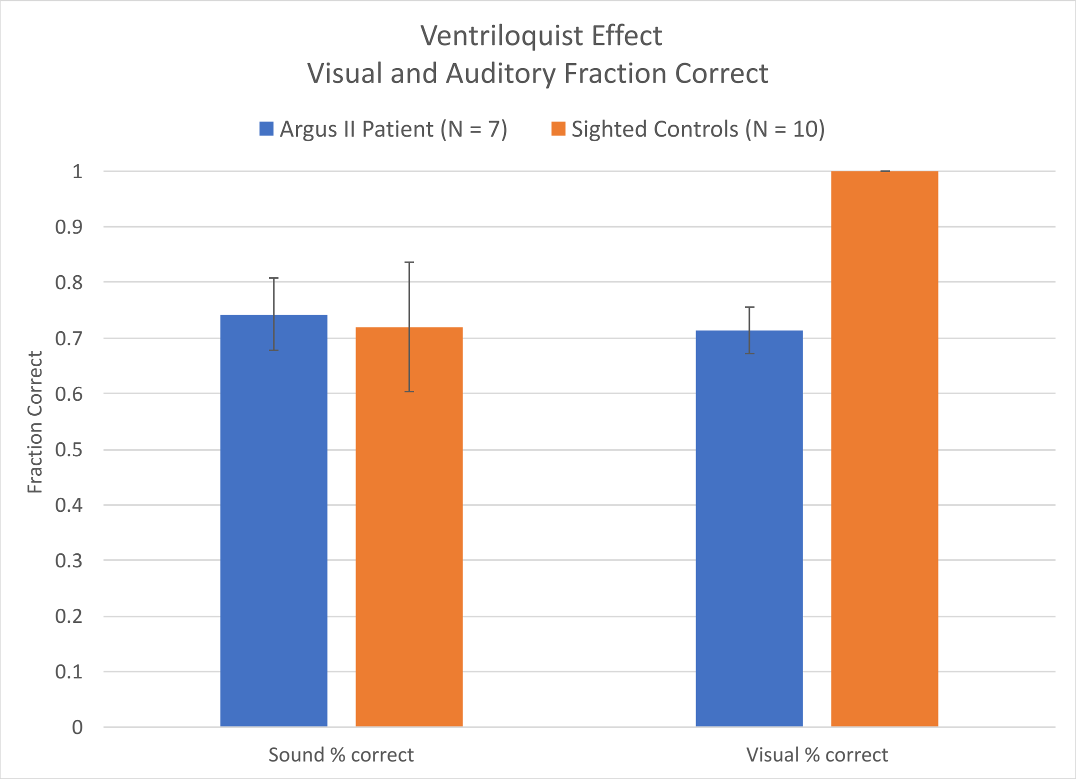


During the ventriloquist experiment, participants performed two baseline tasks: visual alone localization and auditory alone localization. These baseline performance tasks were used to calculate the percent correct at localization by calculating the percentage of trials in which the visual rectangle or auditory beep were localized correctly (within one reporting location out of nine possible; Chance = 1/9).

**Figure S2: Comparison of Ventriloquist Spatial Shift (Visual Angle) Between Argus II Patients and Sighted Controls**

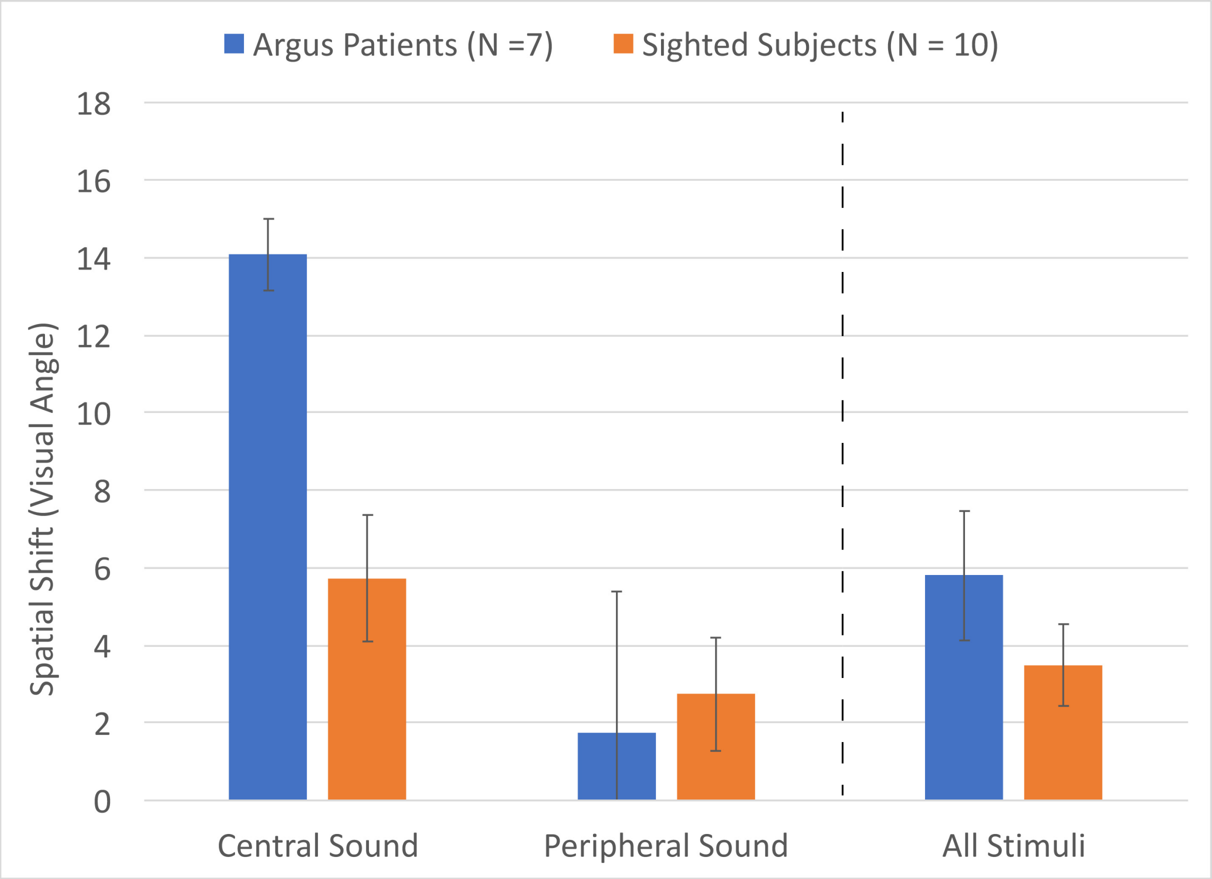


The ventriloquist spatial shift (visual angle) is plotted for both the central sound and the peripheral sound conditions. In this set of plots, the central sound condition represents the cases in which the auditory beep is in the center, and the visual stimulus is either on the left or the right. The peripheral sound condition represents the cases in which the auditory beep is either on the left or the right, and the visual stimulus is in the center. Positive spatial shifts (visual angles) correspond to a reported sound location toward the visual flash, and negative spatial shifts (visual angles) correspond to a reported sound location away from the visual flash. The spatial shifts plotted above represent the spatial shift observed, averaged over the trials for each subject, and averaged over subjects, plotted separately for the Argus II retinal prosthesis patients and the sighted (control) subjects.  The ventriloquist spatial shifts averaged over all of these trials including both participant groups are also shown (All Stimuli).

**Figure S3: Results for Ventriloquist Task with Five Sound Locations in Comparison with the Original Design with Three Sound Locations**

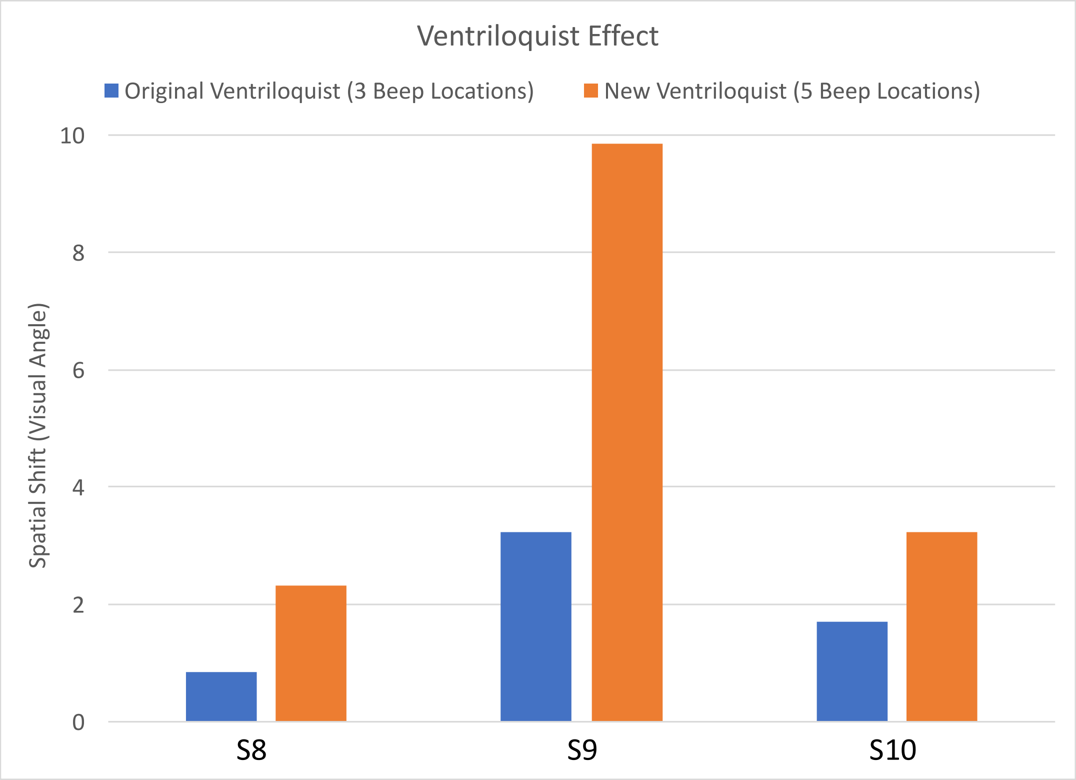


Due to the weaker than expected ventriloquist effect in sighted individuals, an additional experimental design was tested on a subset of participants (Subjects S8, S9, and S10). This second ventriloquist task design used five sound locations and the built-in computer speakers (instead of headphones). The increased auditory ambiguity with the five sound locations increases the influence of vision on audition relative to the three sound location experiment, and resulted in increased angular shifts for all three sighted subjects. This result with a second type of experimental design supports the suggestion in this paper that sighted subjects may exhibit a slightly reduced magnitude of the ventriloquist effect due to their access to external cues that were not available to the Argus II patients.  Increasing the auditory ambiguity for the sighted subjects would tend to offset this empirical advantage in performing the ventriloquist task.

**Figure S4: Schematic Diagram of the Five Sound Location Ventriloquist Experiment**

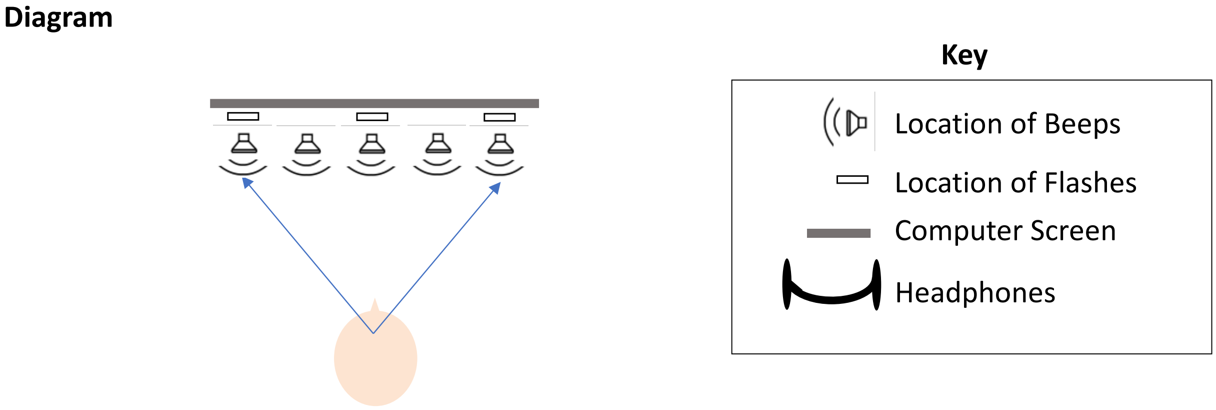


The five sound location ventriloquist task used the built-in computer speakers (instead of headphones) to generate five different auditory beep locations by means of left-right amplitude differences. This figure displays the experimental setup of the five sound location experiment in the same format as that of the three sound location experiment shown in Figure 2. The computer speakers were used to present the auditory stimuli in varying lateral locations by implementing specific variations in the relative auditory amplitudes of the left and right speakers (further details are given in the Supplementary Methods section).

**Figure S5: Double Staircase Ventriloquist Effect Result (*N* = 2)**

A. Double Staircase Result for Subject A7

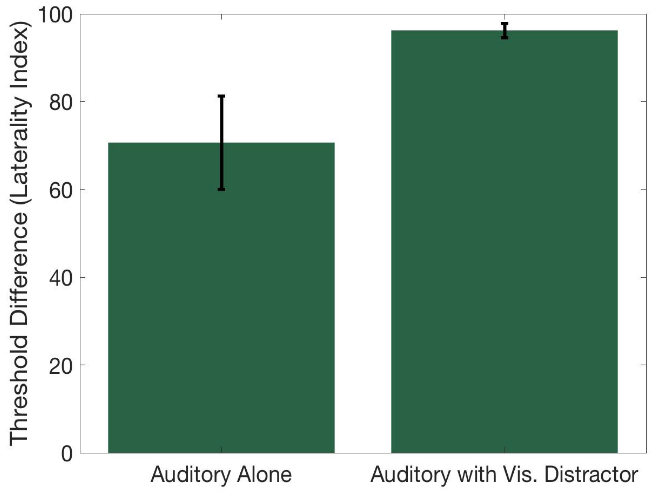


B. Double Staircase Result for Subject A4

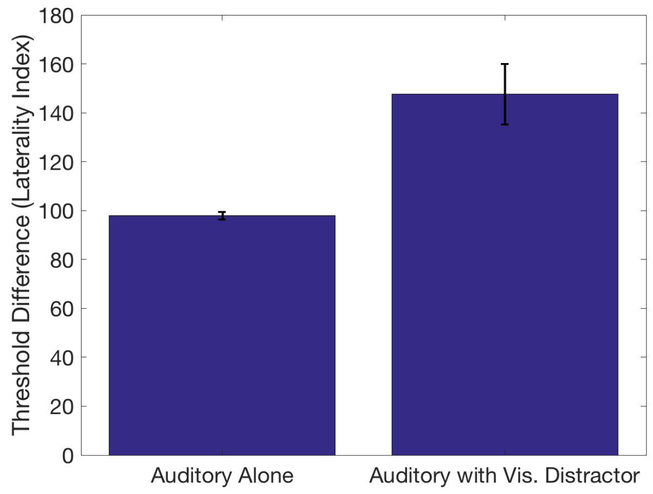


These plots indicate the threshold difference for auditory left-right differentiation (2AFC) *vs.* the type of double staircase tested (auditory alone, or auditory with a visual distractor). The threshold difference (laterality index) represents the difference between two measured thresholds, one for the right staircase (positive valued) and the other for the left staircase (negative valued). This is equivalent to the sum of the absolute values of the two thresholds.  Larger threshold differences (a larger laterality index) indicate that the average left and right thresholds were more lateral (displaced from the center).  The maximum laterality index is 200, which would indicate a +100 laterality index for the right threshold, and a –100 laterality index for the left threshold.  Note:  The right threshold is +100 when the sound balance at the measured threshold condition is set to 100% sound amplitude in the right ear and 0% sound amplitude in the left ear.  Similarly, the left threshold is –100 when the sound balance at the measured threshold condition is set to 0% sound amplitude in the right ear and 100% sound amplitude in the left ear.

**Supplemental Tables**

**Table S1: Argus II Patient Information**

| Initials | Age | Gender | Duration Blind (Years) | Duration with Argus II (Months) | Vision Left | Vision Right | Eye Patch Left | Eye Patch Right |
|---|---|---|---|---|---|---|---|---|
| **A1** | 69 | M | 21 | 28 | LP | LP | Yes | Yes |
| **A2** | 68 | F | 21 | 29.5 | LP | LP | Yes | No |
| **A3** | 69 | F | 16 | 45.5 | No LP | LP | No | Yes |
| **A4** | 69 | M | 24 | 30 | LP | LP | Yes | Yes |
| **A5** | 53 | F | 26 | 27.5 | LP | No LP | Yes | Yes |
| **A6** | 61 | F | 16 | 42 | LP | No LP | Yes | No |
| **A7** | 46 | M | 20 | 19.5 | LP | LP | Yes | Yes |

**M = Male, F = Female, LP = Light Perception, and RP = Retinitis Pigmentosa**

This table details the demographic information and visual perception capabilities of the Argus II patients. The visual disease that caused blindness in each of the patients is Retinitis Pigmentosa (RP). The visual perception for each eye (as reported by the patient) is presented in Columns 6 and 7, with the indication of “LP” for light perception and “No LP” for no light perception. The use of an eye patch during the ventriloquist experiment for each of the patients’ eyes is indicated in Columns 8 and 9, with a Yes for the use of an eye patch and a No for the absence of an eye patch. All participants who reported light perception used an eye patch (with the exception of the right eye for A2, due to a limited supply of eye patches available).

**Table S2: Sighted Control Patient Information**

| Initials | Age | Gender | Glasses or Contacts? | Hearing Loss Left Ear | Hearing Loss Right Ear |
|---|---|---|---|---|---|
| **S1** | 61 | M | Yes | None | None |
| **S2** | 66 | F | Yes | None | None |
| **S3** | 68 | F | Yes | None | None |
| **S4** | 69 | F | Yes | None | None |
| **S5** | 64 | F | No | Mod | Mod |
| **S6** | 58 | F | Yes | None | None |
| **S7** | 63 | F | Yes | None | None |
| **S8** | 69 | F | Yes | None | None |
| **S9** | 55 | M | Yes | None | None |
| **S10** | 62 | M | No | None | None |

**M = Male, F = Female, None = No Hearing Loss, Mild = Mild Hearing Loss, Mod = Moderate Hearing Loss, and Sev = Severe Hearing Loss**

This table details the demographic information and perceptual capabilities (self-reported) of the sighted control participants. The fourth column details if the participants reported wearing glasses or contacts for reading or distance vision. The final two columns detail the self-reported hearing loss of each participant as: No (None), mild (Mild), moderate (Mod), or severe (Sev) hearing loss.

### Questionnaire for the Sighted Subjects

Retinal Prosthesis Control Questionnaire
V07072018

Participant Name:______________________

Email Address: ________________________

Experiment Date:________________

**Basic Information**

Birth Date:_______________

Gender (circle one): Male Female

Wears Glasses or Contacts? (circle one): Yes No

Had Lasik Visual Correction? (circle one): Yes No

Visual Perception Normal (*i.e.* not significantly impaired)? (with glasses if wearing them):

Yes No

Hearing Loss? (circle one): Yes No

If Hearing Loss, Left Ear: No Loss Mild Loss Moderate Loss

Severe Loss

If Hearing Loss, Right Ear: No Loss Mild Loss Moderate Loss

Severe Loss

Cataract Replacement? (circle one): Yes No

If Cataract Replaced (circle one): Replaced in Left Eye Only

Replaced in Right Eye Only Replaced in Both Eyes
